# Pyrethroid Resistance Status and Multiple kdr Mutations (F1534C/L) in Aedes aegypti Populations from Zika-prone Areas in Lamu County, Kenya

**DOI:** 10.64898/2026.05.21.726856

**Authors:** Jane Thiiru, Richard Ochieng, Gladys Kerich, Eunice Achieng, Janet Ambale, Santos Yalwala, Elizabeth Kioko, David Oullo, Charles Waga, Lucy Isabella, Silvanos Oponda, Gerald G. Kellar, Joel Lutomiah, Robert Haynes, John Eads, Fredrick Eyase

## Abstract

*Aedes aegypti* mosquitoes are the primary vectors for dengue, yellow fever, chikungunya and zika virus transmission, posing significant public health risks. In Kenya, these viruses drive disease outbreaks especially, dengue and chikungunya with coastal Kenya being the most affected. Low-level circulation of the Zika virus has been reported in parts of the Kenyan coast, with confirmed cases reported in Lamu County between August to September 2024. Except for yellow fever, there are no approved vaccines or therapeutics, hence vector control remains the most effective means of protection. However, prolonged exposure to insecticides can lead to resistance, threatening these interventions. Therefore, monitoring resistance in mosquito populations is critical to allow for appropriate interventions using effective chemical classes to prevent disease outbreaks. This study aimed to establish the levels of resistance to pyrethroids, and associated markers, among *Ae. aegypti* populations in sections of Lamu County where there had been a recent localized outbreak of Zika. Mosquito eggs were collected from Mkomani, Kashmir, and Kandahar villages in Lamu County, reared and tested for susceptibility to three pyrethroid insecticides (0.75% permethrin, 0.05% Alpha-cypermethrin and 0.05% deltamethrin) using WHO tube assays. Genotyping of knockdown resistance (kdr) mutations L982W, S989P, A1007G, V1016G/I, and F1534C was done using Sanger sequencing. Association between resistant phenotypes and genotypes were inferred. The results varied between the three pyrethroids with high resistance to permethrin observed (6-15% mortality), for deltamethrin mortality ranged between 53-57%, while for alphacypermethrin 88%-99% mortality was observed. Two mutation types and six genotypes were identified at F1534. No other kdr mutations were detected. The CC genotype was significantly associated with 0.75% permethrin resistance in *Ae. aegypti* populations (OR = 2.87, 95% CI: 1.34-6.17, P = 0.0036). The current data show that *Ae. aegypti* from Kandahar, Kashmir and Mkomani villages in Lamu County have developed very high resistance to permethrin, and varying resistance to other pyrethroids, thus threatening pyrethroid-based control strategies in this region, highlighting the need for alternative strategies to control the vector for arboviruses.

## Introduction

*Aedes aegyp*ti is a highly anthropophilic mosquito species, originally native to sub-Saharan Africa (Moore et al., 2013), now widespread in most tropical and subtropical locations across the world (Kraemer et al., 2015; Laporta et al., 2023; Weetman et al., 2018). It is the primary vector to humans for several globally important arboviruses, including dengue, Zika, chikungunya, and yellow fever, which collectively impose a heavy global health burden (Madewell, 2020). Dengue virus is the most prevalent viral infection with half of the world’s population now at risk, with an estimated 100-400 million infections occurring each year, primarily in tropical/subtropical urban areas (Kularatne & Dalugama, 2022; WHO, 2025). Chikungunya virus transmission has affected 2.8 billion people across 104 countries, with 35 million annual infections, and epidemic outbreaks every 6.2 years primarily in Southeast Asia, Africa, and the Americas (Ribeiro dos Santos et al., 2025). Zika virus, originally identified in Uganda in 1947 (Dick et al., 1952), and thereafter, outbreaks in Yap Island (2007) and French Polynesia (2013-2014), spread globally, including a major epidemic in the Americas in 2015-2016 (Kindhauser et al., 2016). Sustained low-level transmission continues in many regions, with ongoing cases reported in the Americas in 2024 and 2025 (Daudt-Lemos et al., 2025; Lin et al., 2025), posing ongoing risks, especially for pregnant women. Yellow fever virus remains a significant public health threat, particularly in tropical regions of Africa and South America, despite becoming less common compared to the past (Angerami et al., 2025; Wang et al., 2025) due to vaccination. Sporadic outbreaks continue to cause significant mortality in regions with low vaccination coverage, such as the Angola and Democratic Republic of Congo outbreaks of 2016 (Ferreira et al., 2022; Otshudiema, 2017; Srivastava et al., 2024; Zhao et al., 2018).

For decades, the coastal region of Kenya, (Kwale, Kilifi, Mombasa, and Lamu), has experienced recurrent disease outbreaks caused by mosquito-borne viruses, establishing the region as a major hotspot for endemic transmission. Low altitude provides a conducive environment for mosquito breeding and high human population composed of both locals and global tourists (Karungu et al., 2019), leading to frequent outbreaks of dengue fever and chikungunya in Mombasa, (Ellis et al., 2015; Eyase et al., 2020; Johnson et al., 1982; Konongoi et al., 2016; Lutomiah et al., 2016), Kilifi, Malindi (Pollett et al., 2021), and Lamu (Wahid et al., 2017). More recently, dengue outbreaks have increased in frequency along the coast, with multiple serotypes circulating endemically (Muthanje et al., 2022). Lamu County is of particular epidemiological importance as the site of Kenya’s first major chikungunya outbreak in 2004-2005 (Wahid et al., 2017). Recently, Lamu County confirmed five cases of Zika virus infection at King Fahd Referral Hospital, underscoring the ongoing threat posed by emerging arboviruses in this locality (Kimita et al., 2025). Despite the increasing burden of arboviruses in the coastal region there is limited documented vector control for *Aedes aegypti* (Lepore et al., 2025).

In the absence of effective vaccines and available treatments, vector control such as environmental management, biological control, and chemical control remains the key strategy for controlling the transmission of arboviruses (Côrtes et al., 2023; Pereira Cabral et al., 2019; WHO, 2024). Globally, pyrethroids, carbamates and organophosphates, remain the most readily used insecticides during periods of high vector density, especially during outbreaks, for timely control of mosquito populations (Dusfour et al., 2019). Kenya has limited documented vector control programs for *Ae. aegypti* (Lepore et al., 2025), however, insecticidal interventions for the control of malaria vectors (including the use of long-lasting insecticidal nets [LLINs], and indoor residual spraying [IRS]) have long been implemented (Ministry of Health, Kenya, 2025). *Ae. aegypti is* a diurnal mosquito, exhibiting peak biting activity during daylight hours, typically in the early morning and late afternoon, which limits the effectiveness of nocturnally targeted interventions such as LLINs and IRS (Chadee, 1988; Jemberie et al., 2025). Methods of applying insecticide targeting *Ae. aegypti* include thermal fogging, ultra-low volume spraying and aerial spraying (Bonds, 2012). Nevertheless, indiscriminate and repetitive use of insecticides in the public and agricultural sector has led to evolution of insecticide resistance (Zulfa et al., 2022).

Pyrethroids, especially permethrin, deltamethrin, and cypermethrin (Tognarelli et al., 2025) are the most widely used insecticides because of their rapid action and low toxicity to mammals (Ahamad & Kumar, 2023). Pyrethroids target voltage-gated sodium channels in mosquito neurons, causing prolonged depolarization and thereby causing rapid paralysis (knockdown), and eventual lethality (Bloomquist, 1996; Dong et al., 2014; Narahashi, 2000). *Ae. aegypti* resistance to pyrethroid primarily involves target-site mutations in the voltage-gated sodium channel gene (VGSC) and enhanced metabolic detoxification by cytochrome P450 monooxygenases, esterases, and glutathione S-transferases (Moyes et al., 2017). To date, over 20 mutations have been identified in the Vgsc of *Aedes aegypti* mosquitoes (Uemura et al., 2024). However, only six have been linked to functional resistance in pyrethroids, namely, F1534C, V1016G, S989P, I1011M, V410L and V253F (Hernandez & Pietrantonio, 2025; Uemura et al., 2024). The F1534C mutation is one of the most common *kdr* mutations in *Ae. aegypti* populations globally, found in all continents, except Australia (Cosme et al., 2020; Kawada et al., 2016; Tognarelli et al., 2025; Uemura et al., 2024) and confers reduced sensitivity to pyrethroids several-fold (Fan & Scott, 2020).

The escalating burden of arboviral diseases in coastal Kenya suggests limited effectiveness of current vector control programs against *Ae. aegypti*. There remains a paucity of data on the resistance status and underlying mechanisms in Kenya and generally in Africa (Moyes et al., 2017). This study, therefore, aimed to evaluate pyrethroids resistance and associated markers, among *Ae. aegypti* populations in sections of Lamu County where there had been a recent localized outbreak of Zika.

## Materials and Methods

### Ethics statement

The study received approval from the Kenya Medical Research Institute (KEMRI) Scientific and Ethics Review Unit under protocols number 3948. It was exempted from review by the Walter Reed Army Institute of Research (WRAIR) Human Subjects Protection Branch, as per WRAIR Policy 25, as there is no human subjects data or samples being analyzed.

### Study Sites

The study was conducted on Lamu Island, which is part of Lamu County, located on the Kenyan coast along the Indian Ocean. Lamu County is one of the six coastal counties in Kenya, situated approximately at latitude 2° 18′ 0″ S and longitude 40° 42′ 0″ E. It covers an area of approximately 6,273 km² and had a population of 143,920 according to the 2019 census. Lamu County is characterized by a tropical coastal climate with average annual temperatures between 26-30°C and two distinct rainy seasons: the long rains from March to May and the short rains from October to December. Lamu County’s economy is based on crop production, livestock, fisheries, tourism, and mining. Within Lamu County, the study focused on Lamu Island villages of Kashmir, Kandahar, and Mkomani based on recent reports of Zika virus cases.

### Mosquito Collection and Rearing

From November 5 to November 10, 2024, ovicups were deployed in both domestic and peri-domestic environments across the three study sites: Mkomani, Kashmir, and Kandahar, to collect *Ae. aegypti* eggs. Ovicups were filled with approximately 300 ml of dechlorinated water and lined with oviposition paper. The ovicups were placed in shaded, sheltered locations such as near water storage containers, and in vegetation around households to maximize egg collection. Each site contained 20 ovicups distributed evenly to capture spatial variation in mosquito breeding activity.

Oviposition papers were collected after 5 days and transported to the Department of Entomology & Vector Borne Infections, Walter Reed Army Institute of Research-Africa insectary in Kisumu. Upon arrival, eggs were air-dried for 2 days to synchronize hatching and then submerged in dechlorinated water to induce larval emergence. Larvae were reared in plastic trays under controlled insectary conditions maintained at 26 ± 2°C temperature, 70-80% relative humidity, and a 12:12 hour light-dark photoperiod to mimic natural environmental conditions and optimize development.

Larvae were fed daily with finely ground fish food (Tetramin®) at a rate adjusted according to larval density and instar stage. Pupae were collected daily and transferred to emergence cages. Adult mosquitoes were maintained on a 10% sucrose solution provided via cotton wool pads, replenished daily. Adult females, aged 3-5 days post-emergence, were used for subsequent bioassays and molecular analyses.

### Insecticide Susceptibility Bioassays

Adult mosquitoes aged 3 to 5 days resulting from the eggs collected in the filial generations (F0 and F1 generation) were preferentially used. The mosquitoes were deprived of sugar for 4 hours before testing to standardize their metabolic state. Susceptibility to three pyrethroid insecticides (0.75% permethrin, 0.05% Alpha-cypermethrin and 0.05% deltamethrin) was assessed using the World Health Organization (WHO) standard tube assay. For each test, batches of 15-25 mosquitoes were exposed to diagnostic concentrations of insecticide-treated paper lining tubes for 1 hour, during which knockdown was monitored at 10, 15, 20, 30, 40, 50, and 60 minutes to record the time course of insecticide effect. After exposure, mosquitoes were transferred to holding tubes and mortality assessed after 24 h. Control groups of mosquitoes were exposed simultaneously to untreated filter papers. Each site was tested in at least three independent replicates.

### Genomic DNA Extraction

Genomic DNA was extracted from individual mosquitoes (both alive and dead) using the Quick-DNA Tissue/Insect Miniprep Kit, following the manufacturer’s protocols with modifications. Briefly, whole mosquitoes were placed directly into 1.5ml Eppendorf Tubes containing 2.4mm Metal Bead Media (Omni International) and 500 μL Bashing BeadTM Buffer. Samples were homogenized by vigorous bead beating using a FastPrep-24 homogenizer. Following lysis, the homogenate was centrifuged at 10,000 × g for 1 minute to pellet debris. Up to 200 µL of the clarified supernatant was transferred to a Zymo-Spin™ III-F Filter column and centrifuged at 8,000 × g for 1 minute. To the filtered lysate, 600 µL of Genomic Lysis Buffer was added and mixed thoroughly. This mixture was then loaded onto a Zymo-Spin™ IICR Column and centrifuged at 10,000 × g for 1 minute to bind the DNA to the silica membrane.The column was washed sequentially with 200 µL DNA Pre-Wash Buffer and 500 µL g-DNA Wash Buffer, each followed by centrifugation at 10,000 × g for 1 minute. Finally, genomic DNA was eluted by adding 60 µL of DNA Elution Buffer directly onto the column matrix and centrifuging at 10,000 × g for 30 seconds.

### Detection of *kdr* Mutations in domain II and domain III of the VGSC gene in *Aedes aegypti*

To identify potential *Kdr* mutations, two fragments of the coding region of the VGSC gene spanning exon 19 to exon 31 (covering the 989, 1011, 1016, 1007, and 1534 coding positions) were amplified from DNA samples. The numbering of amino acid positions was based on the sequence of the house fly (Musca domestica) VGSC gene (GenBank accession number U38813.1). A PCR reaction mix with a total volume of 25 μL was prepared using 12.5 μL of AmpliTaq Gold™ 360 Master Mix, 0.5 μL of forward primer, 0.5 μL of reverse primer, 8.5 μL of nuclease-free water, and 2 μL of DNA template.

For each DNA sample, a multiplex PCR was performed. For domain II (targeting mutations S989P, I1011M/V, L1014F, V1016G/I, and A1007G), amplification was carried out using primers AaSCF1 (5’-AGACAATGTGGATCGCTTCC-3’) and AaSCR4 (5’-GGACGCAATCTGGCTTGTTA-3’). For domain III (detecting the F1534C), primers AaSCF7 (5’-GAGAACTCGCCGATGAACTT-3’) and AaSCR7 (5’-GACGACGAAATCGAACAGGT-3’) were used in the PCR reaction. The PCR parameters were set at 94 °C for 5 min for initial denaturation, 35 cycles of 94 °C for 30 s for denaturation, 57 °C for 30 s for annealing and 72 °C for 1 min extension, followed by final elongation step at 72 °C for 10 min and was set to hold at 4 °C.

### Molecular identification of mosquito species

#### Amplification of COI gene

The mitochondrial cytochrome-c oxidase-I (COI) gene was used to confirm the species identity of mosquitoes. 15 total DNA extracts were used as templates to amplify ∼ 710bp fragments, using the DNA primers pairs, LCOI490 (forward primer): (5’-GGTCAACAAATCATAAAGATATTGG-3’) and HCO2198 (reverse primer): (5’-TAAACTTCAGGGTGACCAAAAAATCA-3’) (Hebert et al., 2003). The total volume PCR reaction mixture was 25 μl, (2 μl of extracted DNA, 12.5 μl of AmpliTaq GoldTM 360 Master mix, 0.5 μl of Forward primer, 0.5 μl of Reverse primer and 8.5 μl of nuclease-free water. The PCR parameters were set at 94 °C for 5 min for initial denaturation, 35 cycles of 94 °C for 30 s for denaturation, 57 °C for 30 s for annealing and 72 °C for 1 min extension, followed by final elongation step at 72 °C for 10 min and was set to hold at 4 °C.

### PCR product visualization

The PCR products were size separated on a 2% agarose gel stained with GelRed Nucleic Acid Stain in water (Biotium, USA). The gel containing size-separated PCR products was visualized by using the Corning® Axygen® Gel Documentation System.

### PCR Product Purification and Sanger Sequencing Reaction Setup

PCR products for *kdr* mutation analysis in the VGSC gene (domains II and III) and for species confirmation via the mitochondrial COI gene were purified using ExoSAP-IT™ PCR Product Cleanup Reagent (Thermo Fisher Scientific, Waltham, MA, USA). Purified PCR products were subjected to Sanger sequencing using BigDye™ Terminator v3.1 Cycle Sequencing Kit (Thermo Fisher Scientific, Waltham, MA, USA).

For each sample, three sequencing reactions were prepared: two targeting domain II mutations (V1016G, S989P, I1011M/V, A1007G, L1014F) using primers AaSCF3 (5’-GTGGAACTTCACCGACTTCA-3’) and AaSCR6 (5’-CGACTTGATCCAGTTTGGAGA-3’), and one targeting domain III mutation (F1534C) using primer AaSCF8 (5’-TAGCTTTCAGCGGCTTCTTC-3’). Additionally, sequencing of the COI gene was performed using primers LCOI490 (5’-GGTCAACAAATCATAAAGATATTGG-3’) and HCO2198 (5’-TAAACTTCAGGGTGACCAAAAAATCA-3’). All primers were used at 4 μM final concentration. Each 10 μL sequencing reaction contained 1 μL BigDye Terminator, 2 μL 5X sequencing buffer, 1 μL primer, 4 μL nuclease-free water, and 4 μL purified PCR product. Thermal cycling was conducted at 95°C for 5 minutes, followed by 30 cycles of 95°C for 15 seconds, 55°C for 30 seconds, and 68°C for 2 minutes 30 seconds, with a final elongation at 68°C for 3 minutes and a hold at 4°C.

Following cycle sequencing, BigDye Terminator products were purified using the BigDye XTerminator™ Purification Kit (Thermo Fisher Scientific, Waltham, MA, USA). For each reaction, 45 μL of SAM Solution and 10 μL of XTerminator Solution were combined to prepare 55 μL of purification master mix, which was added to each well containing the sequencing products. Plates were sealed, vortexed at 1,800 rpm for 20 minutes at room temperature using an IKA™ MS 3 Digital Orbital Shaker (IKA, Staufen, Germany) and centrifuged at 1,000 × g for 2 minutes in a swinging-bucket centrifuge. Purified sequencing products were analyzed by capillary electrophoresis on an ABI 3500/3500xL Genetic Analyzer (Thermo Fisher Scientific, Waltham, MA, USA) using the BDx run module.

### Sanger Sequencing Data Analysis

Chromatogram files (. ab1) were processed using a custom Python v3.13.3 in-house script that trimmed low-quality bases based on a Phred quality score threshold of 20, discarding sequences shorter than 50 bp after trimming to ensure data reliability. Reverse complement sequences for VGSC domain III were generated from the trimmed reads, and all processed sequences were consolidated into a single FASTA file for subsequent alignment and mutation analysis.

The cleaned sequences were aligned using MAFFT v7.511 (Katoh & Standley, 2013) with default parameters. For *Aedes aegypti*, voltage-gated sodium channel (VGSC) gene sequences were aligned and compared against reference VGSC sequences from Musca domestica retrieved from GenBank to identify and confirm the presence of *kdr* mutations at loci V1016G, S989P, I1011M/V, A1007G, and L1014F in domain II, and F1534C in domain III. Mutation sites were manually inspected within the aligned sequences using Jalview v2.11.4.1 (Waterhouse et al., 2009) to ensure accurate mutation calling. For sequences harboring *kdr* mutations, chromatograms were further examined using Chromas v2.6.6 (Technelysium Pty Ltd, South Brisbane, Australia) to validate mutation presence. The quality of the chromatogram was assessed based on clarity, peak height consistency. Double peaks in the sequences were carefully evaluated. DNA sequence polymorphism (DnaSP) (v 6.12.03) (Rozas et al., 2017) was used to define the haplotype phase and compute the genetic parameters including the number of haplotypes (h), the number of polymorphism sites (S), haplotype diversity (Hd) and nucleotide diversity (π). Reference sequences were downloaded from genebank. The Population Analysis with Reticulate Trees (PopArt) software (v1.7) was used to construct TCS haplotype networks to display the putative evolutionary relationships amongst the kdr sequences (Leigh & Bryant, 2015). The maximum likelihood tree for the haplotype was generated and annotated using MEGA v12 software (Kumar et al., 2024).

### Resolving Heterozygous F1534 Codon Ambiguities Using Illumina Amplicon Sequencing with DNA Prep Kit

Sanger sequencing failed to resolve the amino acid codon at the F1534 position in heterozygous samples exhibiting polymorphisms at both the second and third nucleotide positions. To resolve this, Illumina Amplicon Sequencing using DNA Prep Kit was employed to achieve individual allele base-calling. Briefly, *Ae. aegypti* VGSC domain III was amplified as described above using primers AaSCF7 (5’-GAGAACTCGCCGATGAACTT-3’) and AaSCR7 (5’-GACGACGAAATCGAACAGGT-3’). The amplicons were cleaned using AMPure XP beads (Beckman Coulter) following the manufacturer’s instructions to remove primers, nucleotides, and dimers, followed by quantification using a Qubit fluorometer (Thermo Fisher Scientific) with the dsDNA HS Assay Kit. Library preparation was using the Illumina DNA Prep Kit following manufacturer’s instructions. Libraries were quantified, normalized, and pooled before sequencing on an iSeq100 platform in the paired-end mode, with a read length of 2 × 150 bp. Resulting reads were quality-checked using FastQC v0.12.1, trimmed for adapters and low-quality bases with Cutadapt v5.1 and Trimmomatic v0.40 (Bolger et al., 2014), then aligned to the reference domain III VGSC sequence using Bowtie2 v2.5.4 (Langmead & Salzberg, 2012). Aligned reads were processed with Samtools v1.22.1 (Danecek et al., 2021) for sorting and indexing. Variant visualization and confirmation were performed using IGV v2.19.6 (Robinson et al., 2011) to resolve nucleotide ambiguities and accurately determine codons.

### Data Analysis

Mortality was recorded 24 hours post-exposure and analyzed to estimate the percentage mortality per site according to WHO criteria, mortality rates below 90% indicate confirmed resistance, 90-97% possible resistance, and above 98% indicate susceptibility. Permethrin mortality comparisons between filial generations (F_0_ vs. F_1_) within each population were analyzed using Fisher’s exact tests via R’s base fisher.test() function. The association between *kdr* mutations and permethrin resistance was assessed using Fisher’s Exact test coupled with odds ratio (OR) estimation for individual genotypes compared against all other genotypes combined. The Haldane-Anscombe correction was applied where necessary. Statistical significance was determined by p-values, with 95% confidence intervals (CI) calculated to estimate the precision of the ORs. All analyses were conducted using R software (version 4.5.0) and relevant statistical packages.

## Results

### Mosquito Samples and COI Species Confirmation

A total of 860 adult females aged 3–5 days post-emergence (Mkomani, n = 300; Kashmir, n = 300; Kandahar, n = 260) were used for subsequent bioassays and molecular analyses. Molecular confirmation using the mitochondrial COI gene showed identity to reference *Aedes aegypti* sequences.

**Figure 1.**
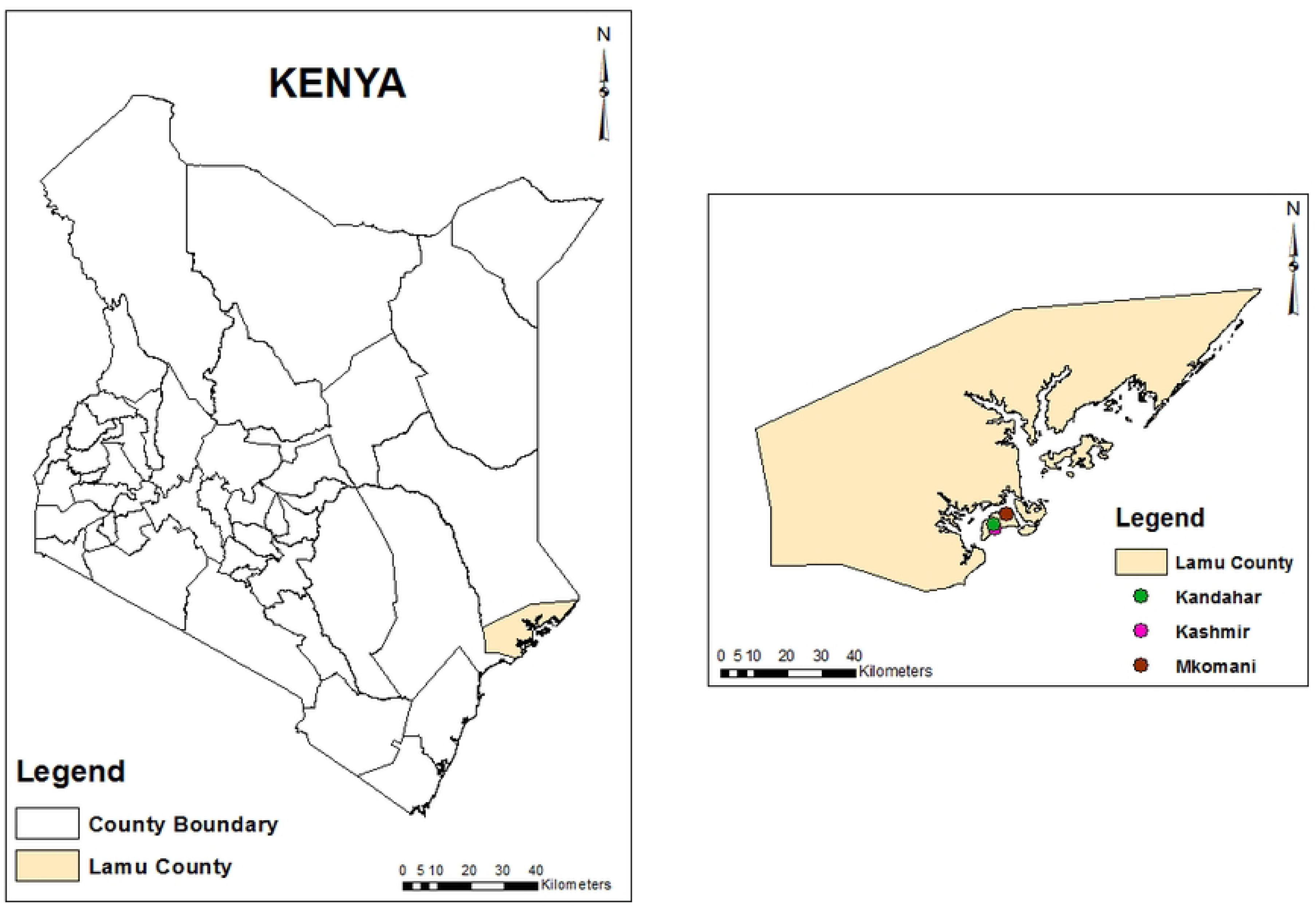
Map of Kenya, Lamu County showing the sampling sites in this study. The map was developed using ArcGIS Software Version 10.2.2 (http://desktop.arcgis.com/en/arcmap) advanced license. https://doi.org/10.1371/journal.pone.0301956.g001

### Bioassay results

Bioassay results showed varying resistance to the three pyrethroids across three *Aedes aegypti* populations (Table 1, Figure 2). Mortality to 0.75% permethrin ranged from 6–15% in F0 and 11–15% in F1 mosquitoes; no significant differences were observed between generations (Fisher’s exact test, P > 0.05), indicating stable resistance across generations. Mortality to 0.05% deltamethrin ranged (53-58%; F1), below the WHO susceptibility threshold of 90%. Alphacypermethrin (0.05%, F1) showed possible resistance in Kashmir-Lamu (88%) and Mkomani-Lamu (93%), but susceptibility in Kandahar-Lamu (99%).

**Figure 2.**
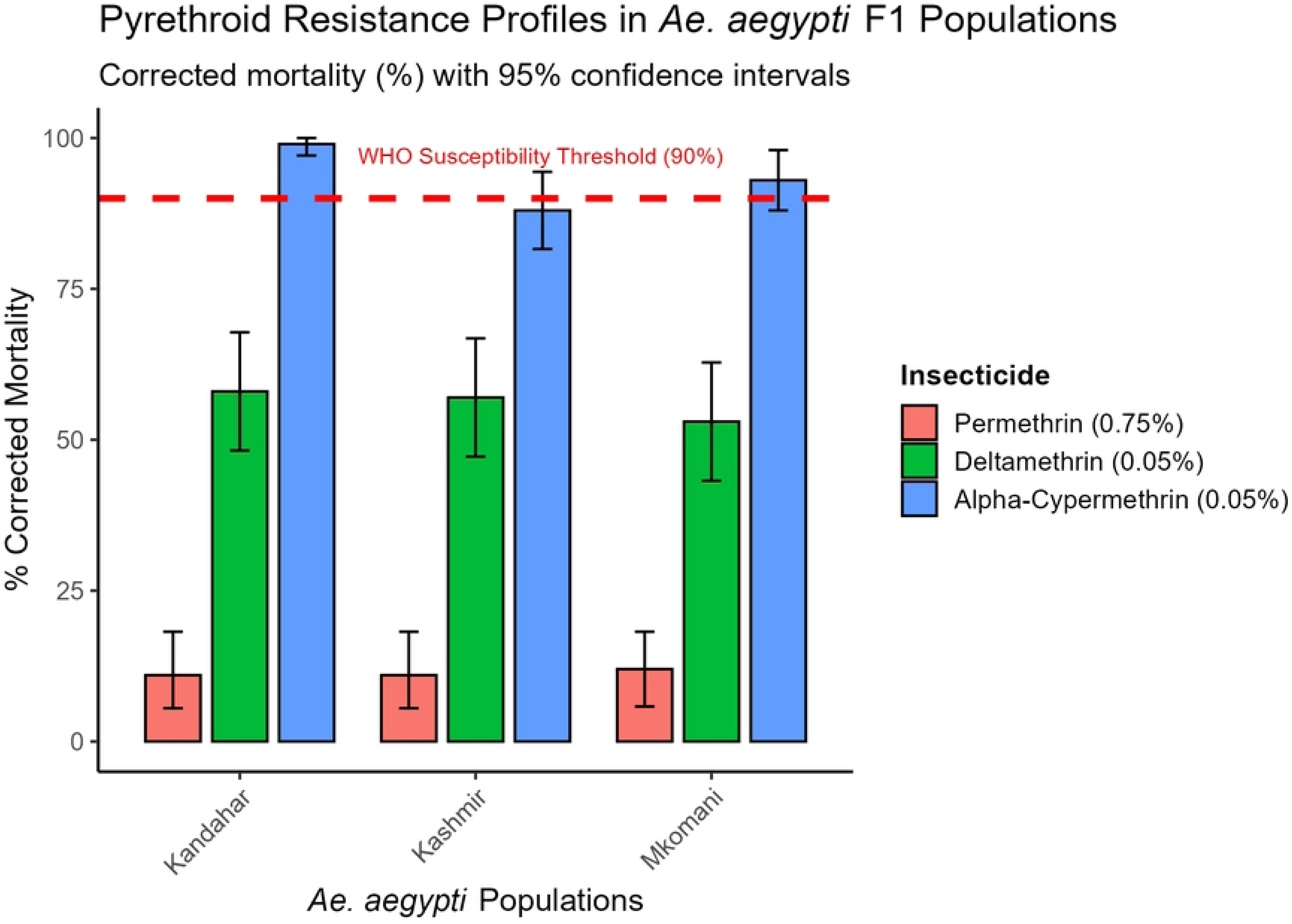
Resistance profiles of permethrin (0.75%), deltamethrin (0.05%) and alpha-cypermethrin (0.05%) in F1 *Ae. aegypti* from three Lamu populations. Bars represent corrected mortality (%); error bars show 95% confidence intervals. Red dashed line marks WHO susceptibility threshold (90% mortality)

**Table 1.** Mortality rates and resistance status of Aedes aegypti populations from Lamu County following exposure to three pyrethroid insecticides using WHO tube assays.

| Insecticide (paper conc) | Population | Date (F Gen) | No. of mosquitoes exposed (n) | % Corrected mortality (95% CI) | Resistance Status |
| --- | --- | --- | --- | --- | --- |
| Permethrin (0.75%) | Kashmir | 11/18/2024 (F0) | 100 | 12% (5.8-18.2) | Confirmed resistance |
|  | Kashmir | 11/20/2024 (F0) | 100 | 13% (6.5-19.5) | Confirmed resistance |
|  | Kashmir | 11/26/2024 (F1) | 100 | 11% (5.5-18.2) | Confirmed resistance |
| Permethrin (0.75%) | Kandahar | 11/20/2024 (F0) | 60 | 8.3% (1.8-18.2) | Confirmed resistance |
|  | Kandahar | 11/25/2024 (F0) | 100 | 6% (1.3-13.2) | Confirmed resistance |
|  | Kandahar | 11/26/2024 (F1) | 100 | 11% (5.5-18.2) | Confirmed resistance |
| Permethrin (0.75%) | Mkomani | 11/18/2024 (F0) | 100 | 15% (8.2-22.7) | Confirmed resistance |
|  | Mkomani | 11/21/2024 (F0) | 100 | 9% (3.6-16.9) | Confirmed resistance |
|  | Mkomani | 11/26/2024 (F1) | 100 | 12% (5.8-18.2) | Confirmed resistance |
| Deltamethrin (0.05%) | Kashmir | 11/27/2024 (F1) | 100 | 57% (47.2-66.8) | Confirmed resistance |
| Deltamethrin (0.05%) | Kandahar | 11/28/2024 (F1) | 100 | 58% (48.2-67.8) | Confirmed resistance |
| Deltamethrin (0.05%) | Mkomani | 11/27/2024 (F1) | 100 | 53% (43.2-62.8) | Confirmed resistance |
| Alpha-cypermethrin (0.05%) | Kashmir | 04/12/2024 (F1) | 100 | 88% (81.6-94.4) | Confirmed resistance |
| Alpha-cypermethrin (0.05%) | Kandahar | 05/12/2024 (F1) | 100 | 99% (97.1-100) | Susceptible |
| Alpha-cypermethrin (0.05%) | Mkomani | 04/12/2024 (F1) | 100 | 93% (88.0-98.0) | Possible resistance |

Percent corrected mortality (with 95% confidence intervals) is shown for each exposure. No mortality was recorded in the controls

### Genotype frequencies

We genotyped VGSC domain III in *Aedes aegypti* exposed to all three pyrethroids: permethrin (n = 302: Mkomani 120, Kashmir 89, Kandahar 93), deltamethrin (n = 164: Mkomani 57, Kashmir 48, Kandahar 59), and Alpha-cypermethrin (n = 172: Mkomani 59, Kashmir 58, Kandahar 55) and domain II only in permethrin-exposed mosquitoes (n = 302: Mkomani 120, Kashmir 89, Kandahar 93). Three alleles were detected: wild-type F1534 (F), and kdr mutants F1534C (C) and F1534L (L), yielding six genotypes; homozygous wild-type F1534 (FF), heterozygotes F1534 (FC) and F1534 (FL), homozygous C1534 (CC), compound heterozygote C1534L (CL), and homozygous L1534 (LL) as shown in Figure 3.

**Figure 3.**
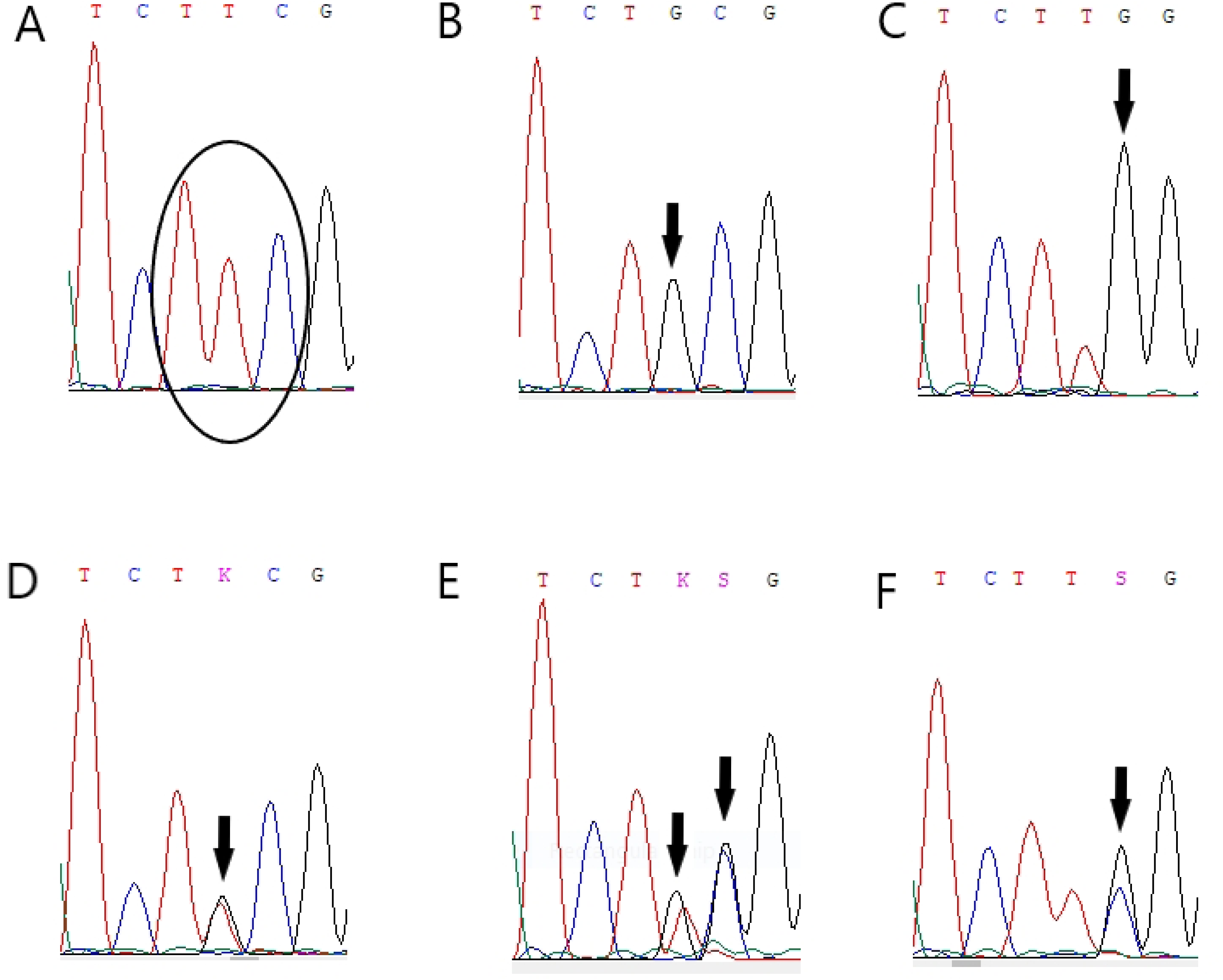
VGSC codon 1534 variants in Aedes aegypti. Sanger sequencing chromatograms showing (A) wild-type, (B) homozygous C1534, (C) homozygous L1534, (D) heterozygote F1534C, (E) compound heterozygote C1534L, and (F) heterozygote F1534L.

**Table 2.**
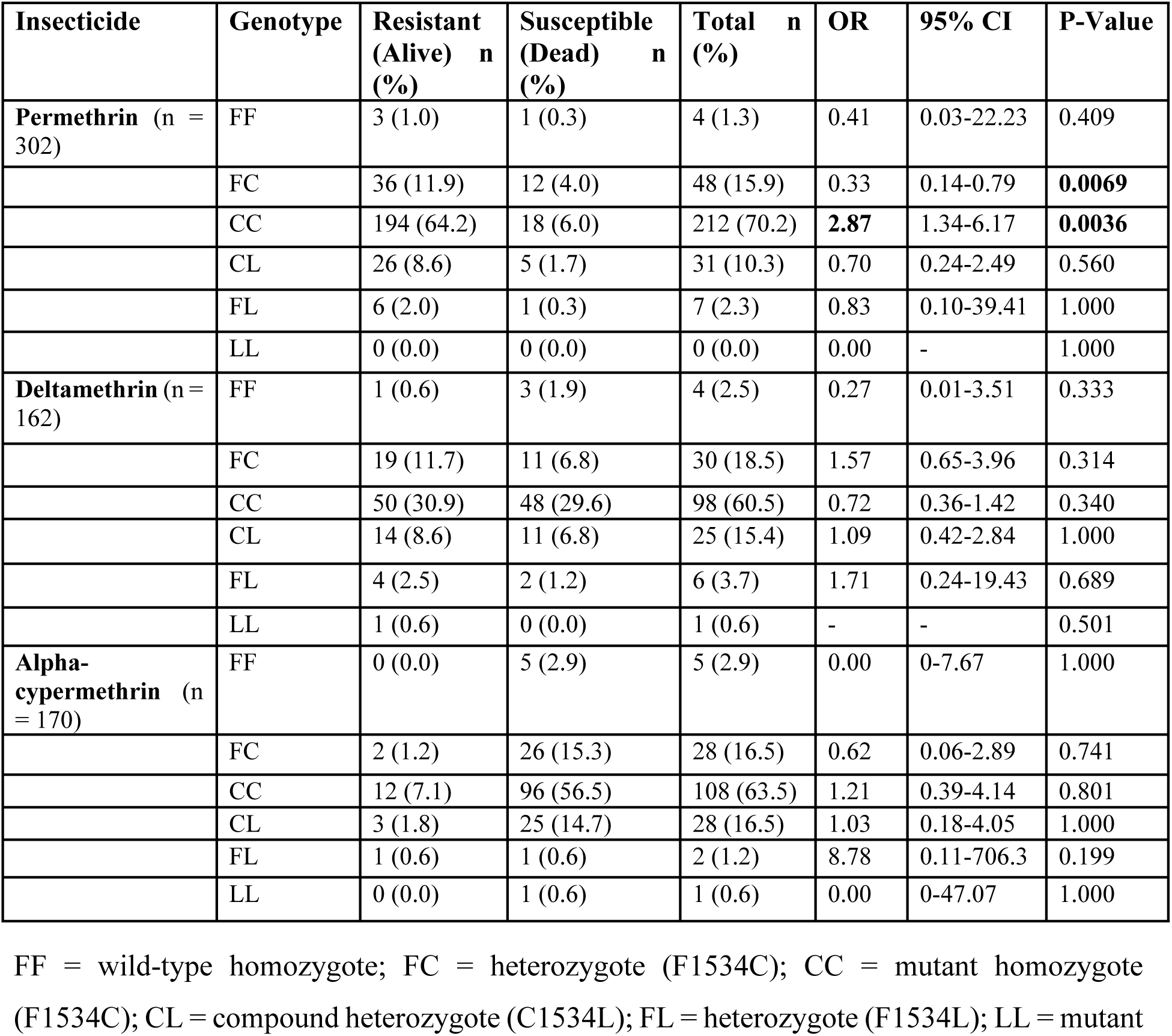
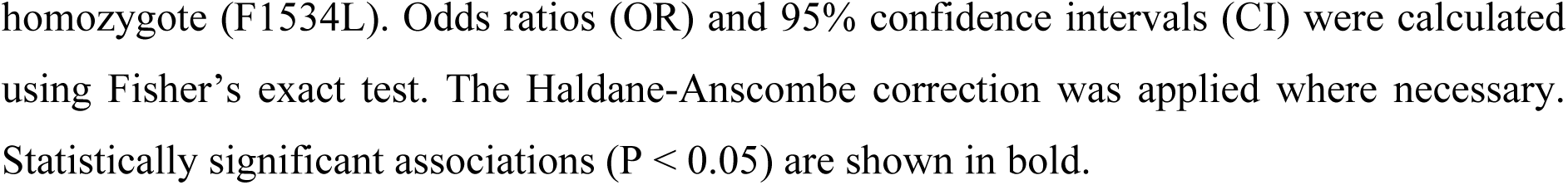
Association between VGSC codon 1534 genotypes and pyrethroid resistance in Aedes aegypti populations from Lamu County, Kenya.

| Insecticide | Genotype | Resistant (Alive) n (%) | Susceptible (Dead) n (%) | Total n (%) | OR | 95% CI | P-Value |
| --- | --- | --- | --- | --- | --- | --- | --- |
| **Permethrin** (n = 302) | FF | 3 (1.0) | 1 (0.3) | 4 (1.3) | 0.41 | 0.03-22.23 | 0.409 |
|  | FC | 36 (11.9) | 12 (4.0) | 48 (15.9) | 0.33 | 0.14-0.79 | **0.0069** |
|  | CC | 194 (64.2) | 18 (6.0) | 212 (70.2) | **2.87** | 1.34-6.17 | **0.0036** |
|  | CL | 26 (8.6) | 5 (1.7) | 31 (10.3) | 0.70 | 0.24-2.49 | 0.560 |
|  | FL | 6 (2.0) | 1 (0.3) | 7 (2.3) | 0.83 | 0.10-39.41 | 1.000 |
|  | LL | 0 (0.0) | 0 (0.0) | 0 (0.0) | 0.00 | - | 1.000 |
| **Deltamethrin** (n = 162) | FF | 1 (0.6) | 3 (1.9) | 4 (2.5) | 0.27 | 0.01-3.51 | 0.333 |
|  | FC | 19 (11.7) | 11 (6.8) | 30 (18.5) | 1.57 | 0.65-3.96 | 0.314 |
|  | CC | 50 (30.9) | 48 (29.6) | 98 (60.5) | 0.72 | 0.36-1.42 | 0.340 |
|  | CL | 14 (8.6) | 11 (6.8) | 25 (15.4) | 1.09 | 0.42-2.84 | 1.000 |
|  | FL | 4 (2.5) | 2 (1.2) | 6 (3.7) | 1.71 | 0.24-19.43 | 0.689 |
|  | LL | 1 (0.6) | 0 (0.0) | 1 (0.6) | - | - | 0.501 |
| **Alpha-cypermethrin** (n = 170) | FF | 0 (0.0) | 5 (2.9) | 5 (2.9) | 0.00 | 0-7.67 | 1.000 |
|  | FC | 2 (1.2) | 26 (15.3) | 28 (16.5) | 0.62 | 0.06-2.89 | 0.741 |
|  | CC | 12 (7.1) | 96 (56.5) | 108 (63.5) | 1.21 | 0.39-4.14 | 0.801 |
|  | CL | 3 (1.8) | 25 (14.7) | 28 (16.5) | 1.03 | 0.18-4.05 | 1.000 |
|  | FL | 1 (0.6) | 1 (0.6) | 2 (1.2) | 8.78 | 0.11-706.3 | 0.199 |
|  | LL | 0 (0.0) | 1 (0.6) | 1 (0.6) | 0.00 | 0-47.07 | 1.000 |

FF = wild-type homozygote; FC = heterozygote (F1534C); CC = mutant homozygote (F1534C); CL = compound heterozygote (C1534L); FL = heterozygote (F1534L); LL = mutant
homozygote (F1534L). Odds ratios (OR) and 95% confidence intervals (CI) were calculated using Fisher's exact test. The Haldane-Anscombe correction was applied where necessary. Statistically significant associations ( $P < 0.05$ ) are shown in bold.

The CC genotype strongly predominated (72.5% Mkomani, 69.7% Kashmir, 53.6% Kandahar). The FC genotype mutation was observed at frequencies ranging from 12.28% in Mkomani to 13.84% in Kashmir and 24.15% in Kandahar as shown in Figure 4. A total of 84 samples had mixed bases at the second and third position of the codon, i.e. with TKS, which could be either heterozygote for Phe/Trp (TTC+ TGG) or Cys/Leu (TGC+ TTG). The former combination was ruled out as sequencing of PCR products using illumina revealed the presence of 2 haplotypes one with TTG (Leu) and another with TGC (Cys) the C1534L(CL) variant. The CL genotype was detected at frequencies of 12.71% in Mkomani, 12.82% in Kashmir, and 14.00% in Kandahar.

**Figure 4.**
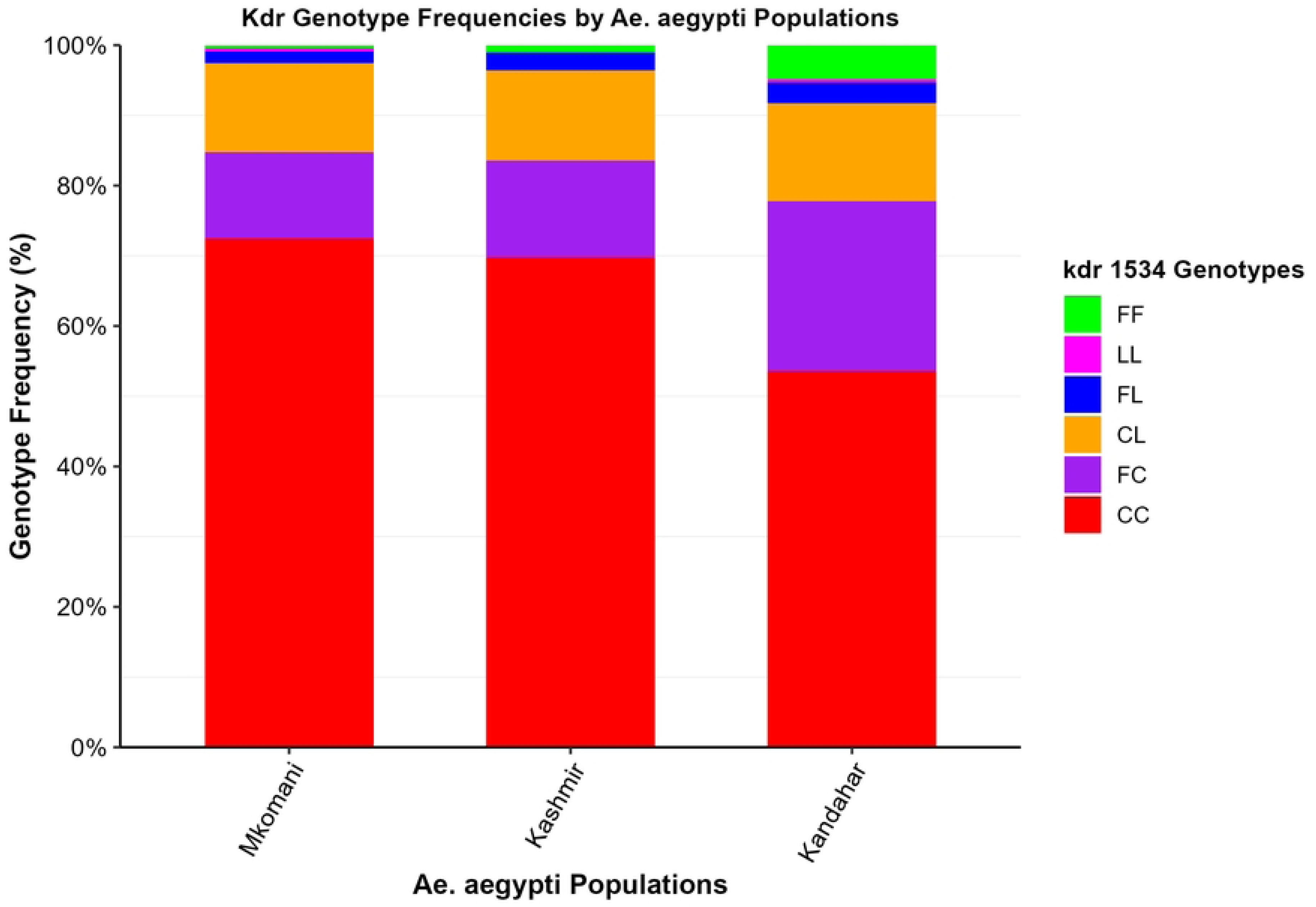
Stacked bar chart of kdr codon 1534 genotype frequencies (%) in *Aedes aegypti* from three Kenyan populations (Mkomani n=236, Kashmir n=195, Kandahar n=207) across pyrethroid exposures. Shows all six VGSC genotypes: FF (F1534, green), LL (L1534, magenta), FL (F1534L, blue), CL (C1534L, orange), FC (F1534C, purple), CC (C1534, red)

The heterozygous FL genotype occurred at low frequencies of 1.69%, 2.56%, and 2.89% in Mkomani, Kashmir, and Kandahar, respectively. The rarest genotype was LL with only two LL homozygotes (0.5%) from Kandahar and Mkomani (0.4%.) Wild-type FF genotype was also infrequent, detected at 0.42% in Mkomani, 1.02% in Kashmir, and 4.83% in Kandahar.

Sequencing of VGSC domain II (covering loci L982W, S989P, V1016G/I, and A1007G) was successfully performed on 302 individuals exposed to permethrin across the three Lamu populations. No mutations associated with pyrethroid resistance were detected at any of these positions.

### Association between kdr mutation at VGSC 1534 codon and pyrethroid resistance

The OR values and 95% CIs were calculated for the six kdr VGSC variants at loci 1534 in *Ae. aegypti* exposed to pyrethroids. The homozygous mutant genotype CC was significantly associated with permethrin resistance (OR = 2.87, 95% CI: 1.34-6.17, P = 0.0036) Table2. Conversely, the heterozygous FC genotype was negatively associated with resistance (OR = 0.33, 95% CI: 0.14-0.79, P = 0.0069). No significant associations were observed for other genotypes to permethrin and no genotype showed a significant association with resistance to deltamethrin or alpha-cypermethrin, indicating that codon 1534 mutations primarily mediate resistance to permethrin in these populations.

### Genetic diversity of VGSC Domain III in *Ae. aegypti*

We analyzed a total of 524 diploid sequences for the VGSC Domain III region associated with permethrin resistance. Within the 354 bp fragment, we identified 11 polymorphic sites that defined 10 distinct haplotypes, as visualized by TCS haplotype network and Maximum likelihood phylogenetic tree in Figure 5 showing low divergence with dominant haplotypes comprising over 80% of sequences. Overall haplotype diversity was low (Hd = 0.277 ± 0.025), accompanied by low nucleotide diversity (π = 0.00285). The most common haplotypes represented 82.7%, 9.3% and 5.2% of sequences, while the remaining haplotypes were rare, each occurring in fewer than 2% of sequences. Neutrality tests showed a negative Fu’s Fs value (Fs = −2.666), although it was not statistically significant (P > 0.10). Tajima’s D was also slightly negative (D = −0.80366, P > 0.10), reflecting low genetic diversity at this locus as shown in Table 3.

**Figure 5.**
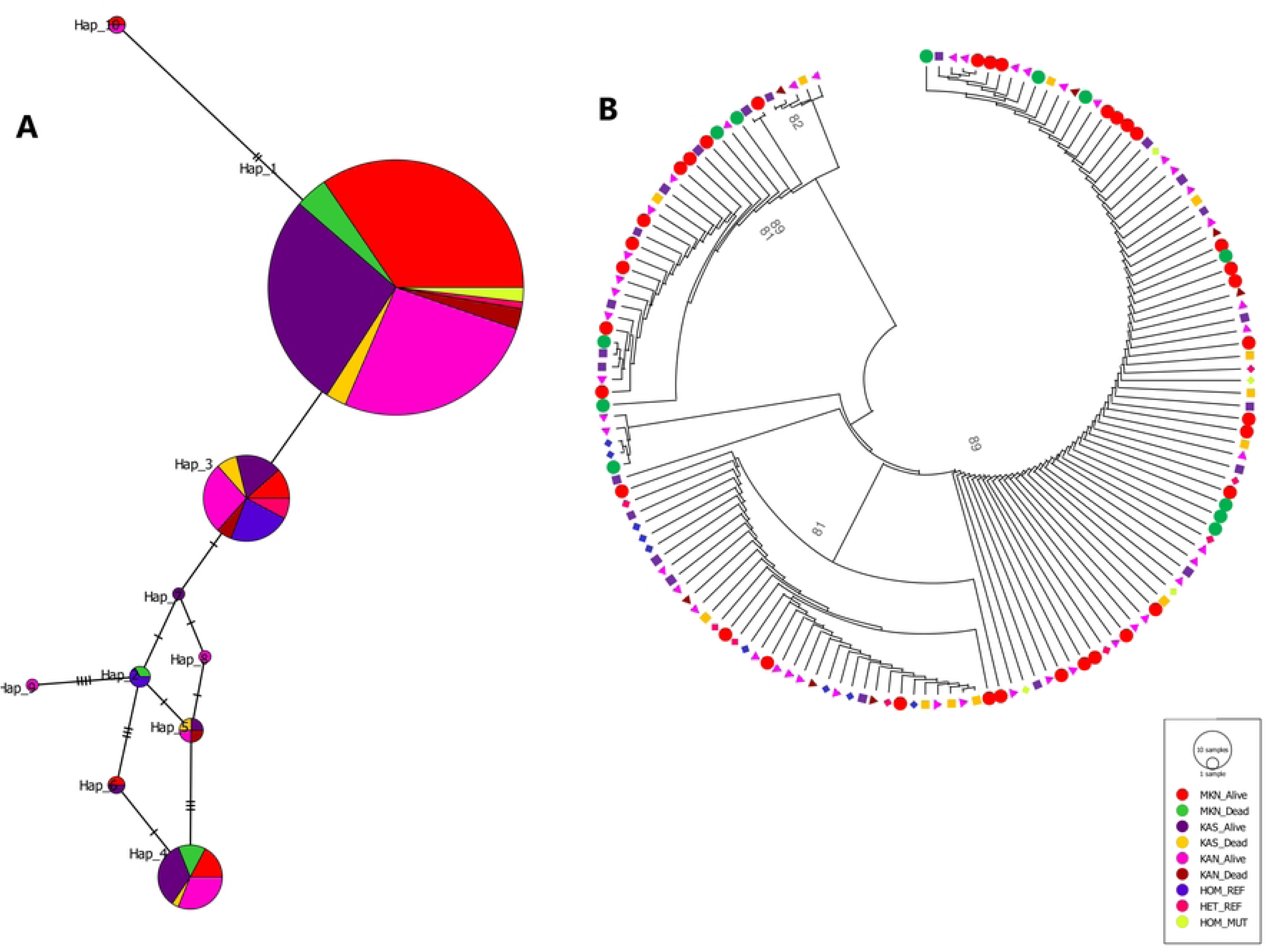
Genetic relationships among VGShC Domain III haplotypes in Aedes aegypti. (A) TCS haplotype network based on a 354 bp alignment, with circle size proportional to haplotype frequency and lines representing mutational steps. (B) Maximum likelihood phylogenetic tree of the 10 haplotypes showing low sequence divergence; bootstrap values (1,000 replicates) are indicated at the nodes.

**Table 3.** Genetic diversity indices of VGSC domain III sequences in Aedes aegypti populations stratified by site and resistance phenotype.

| Samples | 2N | S | $\pi$ | H | $H_d \pm SD$ | Tajima's D | Fu's $F_s$ | p-value |
| --- | --- | --- | --- | --- | --- | --- | --- | --- |
| Kashmir Alive | 148 | 7 | 0.00294 | 7 | 0.258 ± 0.046 | -0.379 | 0.085 | $P > 0.10$ |
| Kashmir Dead | 18 | 7 | 0.00404 | 4 | 0.529 ± 0.118 | -1.007 | 0.722 | $P > 0.10$ |
| Kandahar Alive | 148 | 10 | 0.00331 | 8 | 0.342 ± 0.048 | -0.841 | -1.035 | $P > 0.10$ |
| Kandahar Dead | 16 | 4 | 0.00219 | 3 | 0.425 ± 0.133 | -1.117 | 0.458 | $P > 0.10$ |
| Mkomani Alive | 180 | 9 | 0.00228 | 6 | 0.197 ± 0.039 | -1.113 | -0.576 | $P > 0.10$ |
| Mkomani Dead | 24 | 7 | 0.00619 | 3 | 0.359 ± 0.110 | 0.529 | 3.905 | $P > 0.10$ |
| **Combined Total** | **524** | **11** | **0.00285** | **10** | **0.268 ± 0.025** | **-0.8037** | **-2.666** | **$P > 0.10$** |

### Intron Diversity in VGSC Domain II

Chromatogram analysis and sequence alignment showed large indels in the intron of domain II (236 bp) for all CL variants, however, in addition two types of introns that differed by ∼ 16 bp corresponding to clade A and B introns were observed. Type B intron was the most prevalent associated with the F1534C mutation, while type A intron was observed in samples harbouring heterozygous F1534L and wild type alleles.

## Discussion

Recent detection of a cluster of Zika virus disease cases in Lamu Island (Kimita et al., 2025), alongside the known circulation risk of multiple arboviruses in coastal regions of Kenya (Kirwa et al., 2024), necessitates an effective integrated vector management plan to curb infections. Pyrethroids are the major class of synthetic pesticides that Kenya uses in integrated vector management programs (Karungu et al., 2019). Thus, monitoring efficacy of pyrethroids on *Aedes* vectors is critical.

The present study documents evidence of pyrethroids resistance and multiple *kdr* mutations F1534C/L in three geographically distinct *Aedes aegypti* populations from Lamu, Kenya. To the best of our knowledge, this is the first report documenting pyrethroid resistance and *kdr* mutations in *Ae.aegypti* populations from Kenya. Though the Kenya coastline is considered one of the most endemic regions for mosquito-borne viruses (Karungu et al., 2019), evidence of targeted *Ae Aegypti* control and the effectiveness of control methods remains scarce (Leigh & Bryant, 2015; Lepore et al., 2025), as the numerous anti-malaria campaigns conducted (Ministry of Health, Kenya, 2025), are not effective against day-biting *Ae. aegypti* mosquitoes (Jemberie et al., 2025). The observed permethrin resistance is likely associated with selection pressure arising from the widespread use of pyrethroid insecticides in both integrated vector management programs and agricultural practices (Dusfour et al., 2019).

In this study, mortalities between 6-15% were observed for permethrin, 53-57% for deltamethrin, and 88 - 99% for Alpha-cypermethrin. With respect to permethrin the observed high resistance is consistent with the previous studies carried out in Africa including Côte d’Ivoire (Kadjo et al., 2023), Burkina Faso (Badolo et al., 2019; Yaméogo et al., 2024), Ghana (Kwame Amlalo et al., 2022), Benin (Tokponnon et al., 2024), Congo (Kamgang et al., 2020), Cameroon(Yougang et al., 2020), Nigeria (Mukhtar & Ibrahim, 2022), Niger (Maiga et al., 2024), Mauritania (Haidy Massa et al., 2025) and worldwide (Asgarian et al., 2023). The high mortality observed in both parental (F0) and first filial (F1) generations confirm a stable resistance phenotype that is maintained across generations, with no significant differences observed between them. This sustained resistance poses a critical challenge for vector control efforts relying on pyrethroid insecticides. In addition, the selective pressure for resistance to Alpha-cypermethrin remains relatively low compared to deltamethrin and permethrin.

The dataset reveals the predominance of the F1534C *kdr* mutation across all three populations, with at approximately 95.3% of individuals carrying at least one C allele. This observation aligns closely with the global distribution of this mutation (Cosme et al., 2020; Kawada et al., 2016; Tognarelli et al., 2025; Uemura et al., 2024). Notably, approximately 66 % of the populations are homozygous for the CC mutation, which is significantly associated with pyrethroid resistance (odds ratio = 2.88, p = 0.0036). The intensive use of insecticides exerts a strong selection pressure on mosquitoes, favoring the increase of resistance alleles in natural populations. These findings are consistent with prior reports demonstrating that this mutation reduces the sensitivity of voltage-gated sodium channels to permethrin, thereby conferring resistance (Fan & Scott, 2020; Hu et al., 2011). With the observed deltamethrin, there are conflicting findings in the role of F1534C in deltamethrin resistance (Du et al., 2013; Fan & Scott, 2020) as electrophysiological studies in the Xenopus system showed that this mutation confers reduced VGSC sensitivity to permethrin, but not to deltamethrin (Du et al., 2013).

In addition to F1534C, low-frequency F1534L mutations and CL double heterozygotes were also observed, with CL having a higher frequency. This is the first report of this polymorphism in Kenyan *Ae. aegypti* and suggests ongoing diversification and evolution of resistance mutations at this locus. The F1534L mutation, a recent finding in Indian *Ae. aegypti* populations (Kushwah et al., 2020) associated with permethrin resistance, have since been reported in Myanmar populations (Naw et al., 2022), where it is strongly linked to S989P and V1016G. In this study, no correlation between F1534L heterozygotes and pyrethroid resistance was established.

The very low frequency and near absence of the wild-type FF allele, reflect strong selection pressure and genetic fixation trends toward mutant alleles. The absence of other commonly reported pyrethroid resistance mutations such as V1016G/I, S989P, L982W, and A1007G is interesting given their frequent co-occurrence with F1534C in other *Ae. aegypti* populations (Chen et al., 2021; Dafalla et al., 2019; Hernandez & Pietrantonio, 2025; Kasai et al., 2022; Uemura et al., 2024). This suggests that resistance mechanisms can evolve differently in various geographic regions or under different insecticide selection pressures. It also questions whether F1534C mutation alone is sufficient to produce high-level resistance in these Kenyan populations or more additive kdr mutations not investigated in this study, or that alternate mechanisms contribute to the insecticide resistance. Multiple studies have shown resistance increases additively when the two or more *kdr* mutations exist concurrently (Ishak et al., 2017). Potential role of metabolic resistance mechanisms by detoxification enzyme families, such as mixed-function oxidases and glutathione S-transferases which play significant role in the observed resistant phenotypes (Konkon et al., 2025; Murray & Hribar, 2023; Schluep & Buckner, 2021) cannot be ruled out as synergist assays using metabolic enzyme inhibitors were not investigated.

The results of this study reveal a strong link between specific intron types located between exons 20 and 21 of the gene and F1534C/L mutation. The detection of two types of introns, diverging by approximately 16 bp (Clade A and Clade B), aligns with previous findings identifying two distinct, stably-diverged intron haplotypes in this region (Cosme et al., 2020). The high prevalence of Type B intron in samples harbouring the F1534C mutation is consistent with earlier reports from Asian and Middle Eastern populations, where the F1534C mutation has frequently been associated with a 234 bp (Type B) intron (Chung et al., 2019; Cosme et al., 2020; Kaur et al., 2025) this differs from earlier findings in Africa, where F1534C showed a strong association with the Group A intron but was rarely coupled with the Group B intron (Kawada et al., 2016). Type A intron (approx. 250 bp) in samples harbouring the F1534L mutation and wild-type alleles suggests a separate evolutionary origin for this group.

The dominance of a single resistance-associated haplotype, coupled with low nucleotide and haplotype diversity, suggests that the variation in Domain III is strongly shaped by selective pressure from pyrethroid exposure. This study had several limitations. First, metabolic resistance mechanisms were not directly assessed through synergist bioassays or gene expression analyses. Second, sampling was limited to three sites within Lamu County, which may not represent broader regional patterns. Third, temporal variation in resistance was not evaluated. Future studies incorporating longitudinal sampling and metabolic assays are warranted.

Given the critical public health implications, continued reliance on pyrethroid-based interventions in Lamu County may lead to reduced effectiveness of outbreak response strategies and consequent increases in arboviral disease transmission risk, these results advocate for diversification of control tools, such as the inclusion of non-pyrethroid insecticides or biocontrol methods, and investment in resistance management strategies tailored to regional vector populations.

## Conclusion

This study provides the first definitive evidence of pyrethroid resistance and the presence of multiple kdr mutations, including the predominant F1534C and detection of F1534L, in *Ae. aegypti* populations from Lamu, Kenya. Given the critical implications for arboviral disease control, these results strongly advocate for the diversification of vector control strategies beyond pyrethroids, including other chemical modes of action, biological control, and genetic approaches in controlling *Ae. aegypti*. Furthermore, ongoing surveillance and functional investigations of resistance mechanisms, including metabolic pathways, are essential to inform effective integrated vector management approaches tailored to this high-risk region.

## Acknowledgements

We thank Daniel Ngonga, and Vitalice Opondo for their expert contributions in mosquito sampling. Additionally, we extend our gratitude to Samuel Owaka for his expert map generation.

## Disclaimer

The Material has been reviewed by the Walter Reed Army Institute of Research. There is no objection to its presentation and/or publication. The opinions or assertions contained herein are the private views of the author, and are not to be construed as official, or as reflecting true views of the Department of the Army or the Department of Defense or HJF Medical Research International, Inc. The investigators have adhered to the policies for protection of human subjects as prescribed in AR 70–25.

## References

Ahamad, A., & Kumar, J. (2023). Pyrethroid pesticides: An overview on classification, toxicological assessment and monitoring. Journal of Hazardous Materials Advances, 10, 100284. 10.1016/j.hazadv.2023.100284

Angerami, R. N., Socorro Souza Chaves, T. D., & Rodríguez-Morales, A. J. (2025). Yellow fever outbreaks in South America: Current epidemiology, legacies of the recent past and perspectives for the near future. New Microbes and New Infections, 65, 101580. 10.1016/j.nmni.2025.101580

Asgarian, T. S., Vatandoost, H., Hanafi-Bojd, A. A., & Nikpoor, F. (2023). Worldwide Status of Insecticide Resistance of Aedes aegypti and Ae. Albopictus, Vectors of Arboviruses of Chikungunya, Dengue, Zika and Yellow Fever. Journal of Arthropod-Borne Diseases, 17(1), 1–27. 10.18502/jad.v17i1.13198

Badolo, A., Sombié, A., Pignatelli, P. M., Sanon, A., Yaméogo, F., Wangrawa, D. W., Sanon, A., Kanuka, H., McCall, P. J., & Weetman, D. (2019). Insecticide resistance levels and mechanisms in Aedes aegypti populations in and around Ouagadougou, Burkina Faso. PLOS Neglected Tropical Diseases, 13(5), e0007439. 10.1371/journal.pntd.0007439

Bloomquist, J. R. (1996). Ion Channels as Targets for Insecticides. Annual Review of Entomology, 41(Volume 41, 1996), 163–190. 10.1146/annurev.en.41.010196.001115

Bolger, A. M., Lohse, M., & Usadel, B. (2014). Trimmomatic: A flexible trimmer for Illumina sequence data. Bioinformatics, 30(15), 2114–2120. 10.1093/bioinformatics/btu170

Bonds, J. a. S. (2012). Ultra-low-volume space sprays in mosquito control: A critical review. Medical and Veterinary Entomology, 26(2), 121–130. 10.1111/j.1365-2915.2011.00992.x

Chadee, D. D. (1988). Landing periodicity of the mosquito Aedes aegypti in Trinidad in relation to the timing of insecticidal space-spraying. Medical and Veterinary Entomology, 2(2), 189–192. 10.1111/j.1365-2915.1988.tb00071.x

Chen, H., Zhou, Q., Dong, H., Yuan, H., Bai, J., Gao, J., Tao, F., Ma, H., Li, X., Peng, H., & Ma, Y. (2021). The pattern of kdr mutations correlated with the temperature in field populations of Aedes albopictus in China. Parasites & Vectors, 14(1), 406. 10.1186/s13071-021-04906-z

Chung, H.-H., Cheng, I.-C., Chen, Y.-C., Lin, C., Tomita, T., & Teng, H.-J. (2019). Voltage-gated sodium channel intron polymorphism and four mutations comprise six haplotypes in an Aedes aegypti population in Taiwan. PLOS Neglected Tropical Diseases, 13(3), e0007291. 10.1371/journal.pntd.0007291

Côrtes, N., Lira, A., Prates-Syed, W., Dinis Silva, J., Vuitika, L., Cabral-Miranda, W., Durães-Carvalho, R., Balan, A., Cabral-Marques, O., & Cabral-Miranda, G. (2023). Integrated control strategies for dengue, Zika, and Chikungunya virus infections. Frontiers in Immunology, 14. 10.3389/fimmu.2023.1281667

Cosme, L. V., Gloria-Soria, A., Caccone, A., Powell, J. R., & Martins, A. J. (2020). Evolution of kdr haplotypes in worldwide populations of Aedes aegypti: Independent origins of the F1534C kdr mutation. PLOS Neglected Tropical Diseases, 14(4), e0008219. 10.1371/journal.pntd.0008219

Dafalla, O., Alsheikh, A., Mohammed, W., Shrwani, K., Alsheikh, F., Hobani, Y., & Noureldin, E. (2019). Knockdown resistance mutations contributing to pyrethroid resistance in Aedes aegypti population, Saudi Arabia. Eastern Mediterranean Health Journal = La Revue De Sante De La Mediterranee Orientale = Al-Majallah Al-Sihhiyah Li-Sharq Al-Mutawassit, 25(12), 905–913. 10.26719/emhj.19.081

Danecek, P., Bonfield, J. K., Liddle, J., Marshall, J., Ohan, V., Pollard, M. O., Whitwham, A., Keane, T., McCarthy, S. A., Davies, R. M., & Li, H. (2021). Twelve years of SAMtools and BCFtools. GigaScience, 10(2), giab008. 10.1093/gigascience/giab008

Daudt-Lemos, M., Ramos-Silva, A., Faustino, R., Noronha, T. G. de, Vianna, R. A. de O., Cabral-Castro, M. J., Cardoso, C. A. A., Silva, A. A., & Carvalho, F. R. (2025). Rising Incidence and Spatiotemporal Dynamics of Emerging and Reemerging Arboviruses in Brazil. Viruses, 17(2), 158. 10.3390/v17020158

Dick, G. W. A., Kitchen, S. F., & Haddow, A. J. (1952). Zika Virus (I). Isolations and serological specificity. Transactions of The Royal Society of Tropical Medicine and Hygiene, 46(5), 509–520. 10.1016/0035-9203(52)90042-4

Dong, K., Du, Y., Rinkevich, F., Nomura, Y., Xu, P., Wang, L., Silver, K., & Zhorov, B. S. (2014). Molecular biology of insect sodium channels and pyrethroid resistance. Insect Biochemistry and Molecular Biology, 50, 1–17. 10.1016/j.ibmb.2014.03.012

Du, Y., Nomura, Y., Satar, G., Hu, Z., Nauen, R., He, S. Y., Zhorov, B. S., & Dong, K. (2013). Molecular evidence for dual pyrethroid-receptor sites on a mosquito sodium channel. Proceedings of the National Academy of Sciences, 110(29), 11785–11790. 10.1073/pnas.1305118110

Dusfour, I., Vontas, J., David, J.-P., Weetman, D., Fonseca, D. M., Corbel, V., Raghavendra, K., Coulibaly, M. B., Martins, A. J., Kasai, S., & Chandre, F. (2019). Management of insecticide resistance in the major Aedes vectors of arboviruses: Advances and challenges. PLOS Neglected Tropical Diseases, 13(10), e0007615. 10.1371/journal.pntd.0007615

Ellis, E. M., Neatherlin, J. C., Delorey, M., Ochieng, M., Mohamed, A. H., Mogeni, D. O., Hunsperger, E., Patta, S., Gikunju, S., Waiboic, L., Fields, B., Ofula, V., Konongoi, S. L., Torres-Velasquez, B., Marano, N., Sang, R., Margolis, H. S., Montgomery, J. M., & Tomashek, K. M. (2015). A Household Serosurvey to Estimate the Magnitude of a Dengue Outbreak in Mombasa, Kenya, 2013. PLOS Neglected Tropical Diseases, 9(4), e0003733. 10.1371/journal.pntd.0003733

Eyase, F., Langat, S., Berry, I. M., Mulwa, F., Nyunja, A., Mutisya, J., Owaka, S., Limbaso, S., Ofula, V., Koka, H., Koskei, E., Lutomiah, J., Jarman, R. G., & Sang, R. (2020). Emergence of a novel chikungunya virus strain bearing the E1:V80A substitution, out of the Mombasa, Kenya 2017-2018 outbreak. PLOS ONE, 15(11), e0241754. 10.1371/journal.pone.0241754

Fan, Y., & Scott, J. G. (2020). The F1534C voltage-sensitive sodium channel mutation confers 7- to 16-fold resistance to pyrethroid insecticides in Aedes aegypti. Pest Management Science, 76(6), 2251–2259. 10.1002/ps.5763

Ferreira, F. C. da S. L., Camacho, L. A. B., & Villela, D. A. M. (2022). Occurrence of yellow fever outbreaks in a partially vaccinated population: An analysis of the effective reproduction number. PLOS Neglected Tropical Diseases, 16(9), e0010741. 10.1371/journal.pntd.0010741

Haidy Massa, M., Ould Lemrabott, M. A., Gomez, N., Ould Mohamed Salem Boukhary, A., & Briolant, S. (2025). Insecticide Resistance Status of Aedes aegypti Adults and Larvae in Nouakchott, Mauritania. Insects, 16(3), 288. 10.3390/insects16030288

Hebert, P. D. N., Ratnasingham, S., & DeWaard, J. R. (2003). Barcoding animal life: Cytochrome c oxidase subunit 1 divergences among closely related species. Proceedings of the Royal Society B: Biological Sciences, 270(SUPPL. 1). 10.1098/rsbl.2003.0025

Hernandez, J. R., & Pietrantonio, P. V. (2025). Pyrethroid resistance in Aedes aegypti: Genetic mechanisms worldwide, and recommendations for effective vector control. Parasites & Vectors, 18(1), 409. 10.1186/s13071-025-07010-8

Hu, Z., Du, Y., Nomura, Y., & Dong, K. (2011). A sodium channel mutation identified in *Aedes aegypti* selectively reduces cockroach sodium channel sensitivity to type I, but not type II pyrethroids. Insect Biochemistry and Molecular Biology, 41(1), 9–13. 10.1016/j.ibmb.2010.09.005

Ishak, I. H., Kamgang, B., Ibrahim, S. S., Riveron, J. M., Irving, H., & Wondji, C. S. (2017). Pyrethroid Resistance in Malaysian Populations of Dengue Vector Aedes aegypti Is Mediated by CYP9 Family of Cytochrome P450 Genes. PLOS Neglected Tropical Diseases, 11(1), e0005302. 10.1371/journal.pntd.0005302

Jemberie, W., Dugassa, S., & Animut, A. (2025). Biting Hour and Host Seeking Behavior of Aedes Species in Urban Settings, Metema District, Northwest Ethiopia. Tropical Medicine and Infectious Disease, 10(2), 38. 10.3390/tropicalmed10020038

Johnson, B. K., Ocheng, D., Gichogo, A., Okiro, M., Libondo, D., Kinyanjui, P., & Tukei, P. M. (1982). Epidemic dengue fever caused by dengue type 2 virus in Kenya: Preliminary results of human virological and serological studies. East African Medical Journal, 59(12), 781–784.

Kadjo, Y. M.-A. E., Adja, A. M., Guindo-Coulibaly, N., Zoh, D. D., Traoré, D. F., Assouho, K. F., Sadia-Kacou, M. A. C., Kpan, M. D. S., Yapi, A., & Chandre, F. (2023). Insecticide Resistance and Metabolic Mechanisms in Aedes aegypti from Two Agrosystems (Vegetable and Cotton Crops) in Côte d’Ivoire. Vector-Borne and Zoonotic Diseases, 23(9), 475–485. 10.1089/vbz.2022.0077

Kamgang, B., Wilson-Bahun, T. A., Yougang, A. P., Lenga, A., & Wondji, C. S. (2020). Contrasting resistance patterns to type I and II pyrethroids in two major arbovirus vectors Aedes aegypti and Aedes albopictus in the Republic of the Congo, Central Africa. Infectious Diseases of Poverty, 9(1), 23. 10.1186/s40249-020-0637-2

Karungu, S., Atoni, E., Ogalo, J., Mwaliko, C., Agwanda, B., Yuan, Z., & Hu, X. (2019). Mosquitoes of Etiological Concern in Kenya and Possible Control Strategies. Insects, 10(6), 173. 10.3390/insects10060173

Kasai, S., Itokawa, K., Uemura, N., Takaoka, A., Furutani, S., Maekawa, Y., Kobayashi, D., Imanishi-Kobayashi, N., Amoa-Bosompem, M., Murota, K., Higa, Y., Kawada, H., Minakawa, N., Cuong, T. C., Yen, N. T., Phong, T. V., Keo, S., Kang, K., Miura, K., … Komagata, O. (2022). Discovery of super–insecticide-resistant dengue mosquitoes in Asia: Threats of concomitant knockdown resistance mutations. Science Advances, 8(51), eabq7345. 10.1126/sciadv.abq7345

Katoh, K., & Standley, D. M. (2013). MAFFT Multiple Sequence Alignment Software Version 7: Improvements in Performance and Usability. Molecular Biology and Evolution, 30(4), 772–780. 10.1093/molbev/mst010

Kaur, T., Kushwah, R. S., Pradhan, S., Das, M. K., Kona, M. P., Anushrita, Mittal, R., Weetman, D., Dixit, R., & Singh, O. P. (2025). Knockdown-resistance (kdr) mutations in Indian Aedes aegypti populations: Lack of recombination among haplotypes bearing V1016G, F1534C, and F1534L kdr alleles. PLOS Neglected Tropical Diseases, 19(6), e0013126. 10.1371/journal.pntd.0013126

Kawada, H., Higa, Y., Futami, K., Muranami, Y., Kawashima, E., Osei, J. H. N., Sakyi, K. Y., Dadzie, S., Souza, D. K. de, Appawu, M., Ohta, N., Suzuki, T., & Minakawa, N. (2016). Discovery of Point Mutations in the Voltage-Gated Sodium Channel from African Aedes aegypti Populations: Potential Phylogenetic Reasons for Gene Introgression. PLOS Neglected Tropical Diseases, 10(6), e0004780. 10.1371/journal.pntd.0004780

Kimita, G., Collins, J., Mutai, B., Omuseni, E., Masakhwe, C., Ocholla, S., Lemtudo, A., Awinda, G., Githii, R., Kellar, G., & Waitumbi, J. (2025). Detection of zika virus disease caused by the Asian lineage of the zika virus at Kenya’s Lamu island. International Journal of Infectious Diseases, 154, 107862. 10.1016/j.ijid.2025.107862

Kindhauser, M. K., Allen, T., Frank, V., Santhana, R. S., & Dye, C. (2016). Zika: The origin and spread of a mosquito-borne virus. Bulletin of the World Health Organization, 94(9), 675–686C. 10.2471/BLT.16.171082

Kirwa, L. J., Abkallo, H. M., Nyamota, R., Kiprono, E., Muloi, D., Akoko, J., Lord, J. S., & Bett, B. (2024). Arboviruses in Kenya: A Systematic Review and Meta-analysis of Prevalence (p. 2024.10.17.24315511). medRxiv. 10.1101/2024.10.17.24315511

Konkon, A. K., Aikpon, R., Hoyochi, I., Zoungbédji, D. M., Sovi, A., Salako, A. S., Konkon, C., Dangnon, B., Yahoue, G., Adjovi, R. V., Antoine, L., Ahouandjinou, J., Grace, N. K., Adjottin, B., Baba-Moussa, L., Osse, R., Akogbéto, M., & Padonou, G. G. (2025). Insecticide resistance in Aedes aegypti and Aedes albopictus in southern Benin: Quantification, investigation of kdr mutations, and detection of detoxification enzyme activity. Tropical Medicine and Health, 54(1), 6. 10.1186/s41182-025-00875-6

Konongoi, L., Ofula, V., Nyunja, A., Owaka, S., Koka, H., Makio, A., Koskei, E., Eyase, F., Langat, D., Schoepp, R. J., Rossi, C. A., Njeru, I., Coldren, R., & Sang, R. (2016). Detection of dengue virus serotypes 1, 2 and 3 in selected regions of Kenya: 2011–2014. Virology Journal, 13(1), 182. 10.1186/s12985-016-0641-0

Kraemer, M. U., Sinka, M. E., Duda, K. A., Mylne, A. Q., Shearer, F. M., Barker, C. M., Moore, C. G., Carvalho, R. G., Coelho, G. E., Van Bortel, W., Hendrickx, G., Schaffner, F., Elyazar, I. R., Teng, H.-J., Brady, O. J., Messina, J. P., Pigott, D. M., Scott, T. W., Smith, D. L., … Hay, S. I. (2015). The global distribution of the arbovirus vectors Aedes aegypti and Ae. Albopictus. eLife, 4, e08347. 10.7554/eLife.08347

Kularatne, S. A., & Dalugama, C. (2022). Dengue infection: Global importance, immunopathology and management. Clinical Medicine, 22(1), 9–13. 10.7861/clinmed.2021-0791

Kumar, S., Stecher, G., Suleski, M., Sanderford, M., Sharma, S., & Tamura, K. (2024). MEGA12: Molecular Evolutionary Genetic Analysis Version 12 for Adaptive and Green Computing. Molecular Biology and Evolution, 41, 1–9. 10.1093/molbev/msae263

Kushwah, R. B. S., Kaur, T., Dykes, C. L., Ravi Kumar, H., Kapoor, N., & Singh, O. P. (2020). A new knockdown resistance (kdr) mutation, F1534L, in the voltage-gated sodium channel of Aedes aegypti, co-occurring with F1534C, S989P and V1016G. Parasites & Vectors, 13, 327. 10.1186/s13071-020-04201-3

Kwame Amlalo, G., Akorli, J., Etornam Akyea-Bobi, N., Sowa Akporh, S., Aqua-Baidoo, D., Opoku, M., Frempong, K., Pi-Bansa, S., Boakye, H. A., Joannides, J., Nyarko Osei, J. H., Pwalia, R., Abla Akorli, E., Manu, A., & Dadzie, S. K. (2022). Evidence of High Frequencies of Insecticide Resistance Mutations in Aedes aegypti (Culicidae) Mosquitoes in Urban Accra, Ghana: Implications for Insecticide-based Vector Control of Aedes-borne Arboviral Diseases. Journal of Medical Entomology, 59(6), 2090–2101. 10.1093/jme/tjac120

Langmead, B., & Salzberg, S. L. (2012). Fast gapped-read alignment with Bowtie 2. Nature Methods, 9(4), 357–359. 10.1038/nmeth.1923

Laporta, G. Z., Potter, A. M., Oliveira, J. F. A., Bourke, B. P., Pecor, D. B., & Linton, Y.-M. (2023). Global Distribution of Aedes aegypti and Aedes albopictus in a Climate Change Scenario of Regional Rivalry. Insects, 14(1), 49. 10.3390/insects14010049

Leigh, J. W., & Bryant, D. (2015). POPART: Full-feature software for haplotype network construction. Methods in Ecology and Evolution, 6(9), 1110–1116. 10.1111/2041-210X.12410

Lepore, L., Vanlerberghe, V., Verdonck, K., Metelo, E., Diallo, M., & Bortel, W. V. (2025). Vector control for Aedes aegypti and Aedes albopictus mosquitoes implemented in the field in sub-Saharan Africa: A scoping review. PLOS Neglected Tropical Diseases, 19(7), e0013203. 10.1371/journal.pntd.0013203

Lin, L., Guo, X., Wu, Y., Sun, Y., Liu, D., Luo, Y., Xiang, Q., Li, T., Li, G., Yu, W., & Gu, D. (2025). Global Status of Infectious Diseases from January to June of 2025. Zoonoses, 5, 967. 10.15212/ZOONOSES-2025-1002

Lutomiah, J., Barrera, R., Makio, A., Mutisya, J., Koka, H., Owaka, S., Koskei, E., Nyunja, A., Eyase, F., Coldren, R., & Sang, R. (2016). Dengue Outbreak in Mombasa City, Kenya, 2013–2014: Entomologic Investigations. PLOS Neglected Tropical Diseases, 10(10), e0004981. 10.1371/journal.pntd.0004981

Madewell, Z. J. (2020). Arboviruses and Their Vectors. Southern Medical Journal, 113(10), 520–523. 10.14423/SMJ.0000000000001152

Maiga, A.-A., Sombié, A., Zanré, N., Yaméogo, F., Iro, S., Testa, J., Sanon, A., Koita, O., Kanuka, H., McCall, P. J., Weetman, D., & Badolo, A. (2024). First report of V1016I, F1534C and V410L kdr mutations associated with pyrethroid resistance in Aedes aegypti populations from Niamey, Niger. PLOS ONE, 19(5), e0304550. 10.1371/journal.pone.0304550

Ministry of Health, Kenya. (2025). Kenya Malaria Strategy 2023-2027. https://www.afro.who.int/sites/default/files/2025-03/Kenya%20Malaria%20Strategy%20%7C%202023-2027.pdf

Moore, M., Sylla, M., Goss, L., Burugu, M. W., Sang, R., Kamau, L. W., Kenya, E. U., Bosio, C., Munoz, M. de L., Sharakova, M., & Black, W. C. (2013). Dual African Origins of Global Aedes aegypti s.l. Populations Revealed by Mitochondrial DNA. PLOS Neglected Tropical Diseases, 7(4), e2175. 10.1371/journal.pntd.0002175

Moyes, C. L., Vontas, J., Martins, A. J., Ng, L. C., Koou, S. Y., Dusfour, I., Raghavendra, K., Pinto, J., Corbel, V., David, J.-P., & Weetman, D. (2017). Contemporary status of insecticide resistance in the major Aedes vectors of arboviruses infecting humans. PLOS Neglected Tropical Diseases, 11(7), e0005625. 10.1371/journal.pntd.0005625

Mukhtar, M. M., & Ibrahim, S. S. (2022). Temporal Evaluation of Insecticide Resistance in Populations of the Major Arboviral Vector Aedes Aegypti from Northern Nigeria. Insects, 13(2), 187. 10.3390/insects13020187

Murray, H. L., & Hribar, L. J. (2023). Resistance and inhibitor testing on Aedes aegypti (Linnaeus) (Culicidae: Diptera) populations in the Florida Keys. Journal of Vector Ecology, 49(1), 53–63. 10.52707/1081-1710-49.1.53

Muthanje, E. M., Kimita, G., Nyataya, J., Njue, W., Mulili, C., Mugweru, J., Mutai, B., Kituyi, S. N., & Waitumbi, J. (2022). March 2019 dengue fever outbreak at the Kenyan south coast involving dengue virus serotype 3, genotypes III and V. PLOS Global Public Health, 2(3), e0000122. 10.1371/journal.pgph.0000122

Narahashi, T. (2000). Neuroreceptors and ion channels as the basis for drug action: Past, present, and future. The Journal of Pharmacology and Experimental Therapeutics, 294(1), 1–26.

Naw, H., Võ, T. C., Lê, H. G., Kang, J.-M., Mya, Y. Y., Myint, M. K., Kim, T.-S., Shin, H.-J., & Na, B.-K. (2022). Knockdown Resistance Mutations in the Voltage-Gated Sodium Channel of Aedes aegypti (Diptera: Culicidae) in Myanmar. Insects, 13(4), 322. 10.3390/insects13040322

Otshudiema, J. O. (2017). Yellow Fever Outbreak—Kongo Central Province, Democratic Republic of the Congo, August 2016. MMWR. Morbidity and Mortality Weekly Report, 66. 10.15585/mmwr.mm6612a5

Pereira Cabral, B., da Graça Derengowski Fonseca, M., & Mota, F. B. (2019). Long term prevention and vector control of arboviral diseases: What does the future hold? International Journal of Infectious Diseases, 89, 169–174. 10.1016/j.ijid.2019.10.002

Pollett, S., Gathii, K., Figueroa, K., Rutvisuttinunt, W., Srikanth, A., Nyataya, J., Mutai, B. K., Awinda, G., Jarman, R. G., Berry, I. M., & Waitumbi, J. N. (2021). The evolution of dengue-2 viruses in Malindi, Kenya and greater East Africa: Epidemiological and immunological implications. Infection, Genetics and Evolution, 90, 104617. 10.1016/j.meegid.2020.104617

Ribeiro dos Santos, G., Jawed, F., Mukandavire, C., Deol, A., Scarponi, D., Mboera, L. E. G., Seruyange, E., Poirier, M. J. P., Bosomprah, S., Udeze, A. O., Dellagi, K., Hozé, N., Chilongola, J., Nasrallah, G. K., Cauchemez, S., & Salje, H. (2025). Global burden of chikungunya virus infections and the potential benefit of vaccination campaigns. Nature Medicine, 31(7), 2342–2349. 10.1038/s41591-025-03703-w

Robinson, J. T., Thorvaldsdóttir, H., Winckler, W., Guttman, M., Lander, E. S., Getz, G., & Mesirov, J. P. (2011). Integrative genomics viewer. Nature Biotechnology, 29(1), 24–26. 10.1038/nbt.1754

Rozas, J., Ferrer-Mata, A., Sánchez-DelBarrio, J. C., Guirao-Rico, S., Librado, P., Ramos-Onsins, S. E., & Sánchez-Gracia, A. (2017). DnaSP 6: DNA Sequence Polymorphism Analysis of Large Data Sets. Molecular Biology and Evolution, 34(12), 3299–3302. 10.1093/molbev/msx248

Schluep, S. M., & Buckner, E. A. (2021). Metabolic Resistance in Permethrin-Resistant Florida Aedes aegypti (Diptera: Culicidae). Insects, 12(10). 10.3390/insects12100866

Srivastava, S., Dhoundiyal, S., Kumar, S., Kaur, A., Khatib, M. N., Gaidhane, S., Zahiruddin, Q. S., Mohanty, A., Henao-Martinez, A. F., Krsak, M., Rodriguez-Morales, A. J., Montenegro-Idrogo, J. J., Bonilla-Aldana, D. K., & Sah, R. (2024). Yellow Fever: Global Impact, Epidemiology, Pathogenesis, and Integrated Prevention Approaches. Le Infezioni in Medicina, 32(4), 434–450. 10.53854/liim-3204-3

Tognarelli, J., Moya, P. R., González, C. R., & Collao-Ferrada, X. (2025). Global distribution and impact of knockdown resistance mutations in Aedes aegypti on pyrethroid resistance. Parasites & Vectors, 18, 382. 10.1186/s13071-025-06817-9

Tokponnon, T. F., Ossè, R., Zoulkifilou, S. D., Amos, G., Festus, H., Idayath, G., Sidick, A., Messenger, L. A., & Akogbeto, M. (2024). Insecticide Resistance in Aedes aegypti Mosquitoes: Possible Detection of kdr F1534C, S989P, and V1016G Triple Mutation in Benin, West Africa. Insects, 15(4), 295. 10.3390/insects15040295

Uemura, N., Itokawa, K., Komagata, O., & Kasai, S. (2024). Recent advances in the study of knockdown resistance mutations in *Aedes* mosquitoes with a focus on several remarkable mutations. Current Opinion in Insect Science, 63, 101178. 10.1016/j.cois.2024.101178

Wahid, B., Ali, A., Rafique, S., & Idrees, M. (2017). Global expansion of chikungunya virus: Mapping the 64-year history. International Journal of Infectious Diseases, 58, 69–76. 10.1016/j.ijid.2017.03.006

Wang, X., Li, B., He, B., Yan, X., Huang, L., Li, J., Lai, R., Lai, M., Xie, H., Mo, Q., & Chen, L. (2025). The Incidence and Trends of Yellow Fever from 1990 to 2021 in Major Endemic Regions: A Systematic Analysis Based on the 2021 Global Burden of Disease Study. Pathogens, 14(6), 594. 10.3390/pathogens14060594

Waterhouse, A. M., Procter, J. B., Martin, D. M. A., Clamp, M., & Barton, G. J. (2009). Jalview Version 2—A multiple sequence alignment editor and analysis workbench. Bioinformatics, 25(9), 1189–1191. 10.1093/bioinformatics/btp033

Weetman, D., Kamgang, B., Badolo, A., Moyes, C. L., Shearer, F. M., Coulibaly, M., Pinto, J., Lambrechts, L., & McCall, P. J. (2018). Aedes Mosquitoes and Aedes-Borne Arboviruses in Africa: Current and Future Threats. International Journal of Environmental Research and Public Health, 15(2), 220. 10.3390/ijerph15020220

WHO. (2024). WHO launches global strategic plan to fight rising dengue and other Aedes-borne arboviral diseases. https://www.who.int/news/item/03-10-2024-who-launches-global-strategic-plan-to-fight-rising-dengue-and-other-aedes-borne-arboviral-diseases

WHO. (2025, August 21). Dengue,. https://www.who.int/news-room/fact-sheets/detail/dengue-and-severe-dengue

Yaméogo, F., Sombié, A., Oté, M., Saiki, E., Sakurai, T., Wangrawa, D. W., McCall, P. J., Weetman, D., Kanuka, H., & Badolo, A. (2024). Three years of insecticide resistance evolution and associated mechanisms in Aedes aegypti populations of Ouagadougou, Burkina Faso. PLOS Neglected Tropical Diseases, 18(12), e0012138. 10.1371/journal.pntd.0012138

Yougang, A. P., Kamgang, B., Bahun, T. A. W., Tedjou, A. N., Nguiffo-Nguete, D., Njiokou, F., & Wondji, C. S. (2020). First detection of F1534C knockdown resistance mutation in Aedes aegypti (Diptera: Culicidae) from Cameroon. Infectious Diseases of Poverty, 9, 152. 10.1186/s40249-020-00769-1

Zhao, S., Stone, L., Gao, D., & He, D. (2018). Modelling the large-scale yellow fever outbreak in Luanda, Angola, and the impact of vaccination. PLOS Neglected Tropical Diseases, 12(1), e0006158. 10.1371/journal.pntd.0006158

Zulfa, R., Lo, W.-C., Cheng, P.-C., Martini, M., & Chuang, T.-W. (2022). Updating the Insecticide Resistance Status of Aedes aegypti and Aedes albopictus in Asia: A Systematic Review and Meta-Analysis. Tropical Medicine and Infectious Disease, 7(10), 306. 10.3390/tropicalmed7100306

